# A mosaic of whole-body representations in human motor cortex

**DOI:** 10.1101/2024.09.14.613041

**Authors:** Darrel R. Deo, Elizaveta V. Okorokova, Anna L. Pritchard, Nick V. Hahn, Nicholas S. Card, Samuel R. Nason-Tomaszewski, Justin Jude, Thomas Hosman, Eun Young Choi, Deqiang Qiu, Yuguang Meng, Maitreyee Wairagkar, Claire Nicolas, Foram B. Kamdar, Carrina Iacobacci, Alexander Acosta, Leigh R. Hochberg, Sydney S. Cash, Ziv M. Williams, Daniel B. Rubin, David M. Brandman, Sergey D. Stavisky, Nicholas AuYong, Chethan Pandarinath, John E. Downey, Sliman J. Bensmaia, Jaimie M. Henderson, Francis R. Willett

## Abstract

Understanding how the body is represented in motor cortex is key to understanding how the brain controls movement. The precentral gyrus (PCG) has long been thought to contain largely distinct regions for the arm, leg and face (represented by the “motor homunculus”). However, mounting evidence has begun to reveal a more intermixed, interrelated and broadly tuned motor map. Here, we revisit the motor homunculus using microelectrode array recordings from 20 arrays that broadly sample PCG across 8 individuals, creating a comprehensive map of human motor cortex at single neuron resolution. We found whole-body representations throughout all sampled points of PCG, contradicting traditional leg/arm/face boundaries. We also found two speech-preferential areas with a broadly tuned, orofacial-dominant area in between them, previously unaccounted for by the homunculus. Throughout PCG, movement representations of the four limbs were interlinked, with homologous movements of different limbs (e.g., toe curl and hand close) having correlated representations. Our findings indicate that, while the classic homunculus aligns with each area’s preferred body region at a coarse level, at a finer scale, PCG may be better described as a mosaic of functional zones, each with its own whole-body representation.

## Introduction

Nearly a century ago, Penfield and colleagues pioneered the motor homunculus, a topographical map of the human precentral gyrus (PCG), by electrically stimulating the cortical surface and observing which body parts moved in response^1,2^. Although Penfield found large variability across people in the cortical location of each body part, the relative ordering of body parts was thought to be “almost invariable”^1^, with a medial-to-lateral arrangement of leg, arm and orofacial movements. To date, various functional magnetic resonance imaging^3,4^ (fMRI), magnetoencephalography^5^ (MEG), and electrocorticographic^6–9^ (ECoG) studies have largely supported Penfield’s original findings (albeit with occasional exceptions), demonstrating a “macroscopic somatotopy” of mostly separate arm, leg and face areas, with mixing of nearby body parts within those regions (e.g., wrist and fingers may overlap within arm area).

However, these recording modalities average the activity across populations of neurons and cannot resolve detailed representations at the single-neuron level. In prior work using microelectrode arrays capable of recording brain activity at single-neuron resolution, we have shown that a small, anatomically distinct area of cortex in the dorsal PCG (referred to as the ’hand knob’^10^) contained intermixed representations of the entire body^11,12^ (including all four limbs and head and face movements). More recent fMRI^13^ and sEEG^14^ studies have also identified whole-body representations in motor cortex concentrated in specific regions.

Here, we revisit the motor representation of the whole body across a wide span of PCG using recordings from 20 microelectrode arrays in 8 human participants (Fig. 1a) enrolled in brain-computer interface clinical trials. These participants had either spinal cord injury, amyotrophic lateral sclerosis, or brainstem stroke. These recordings collectively sample the length of PCG spanning from the canonical arm area (near the superior frontal sulcus) to the canonical tongue/throat area (near the sylvian fissure), and together constitute the first comprehensive motor map of human PCG at single neuron resolution.

**Fig. 1.**
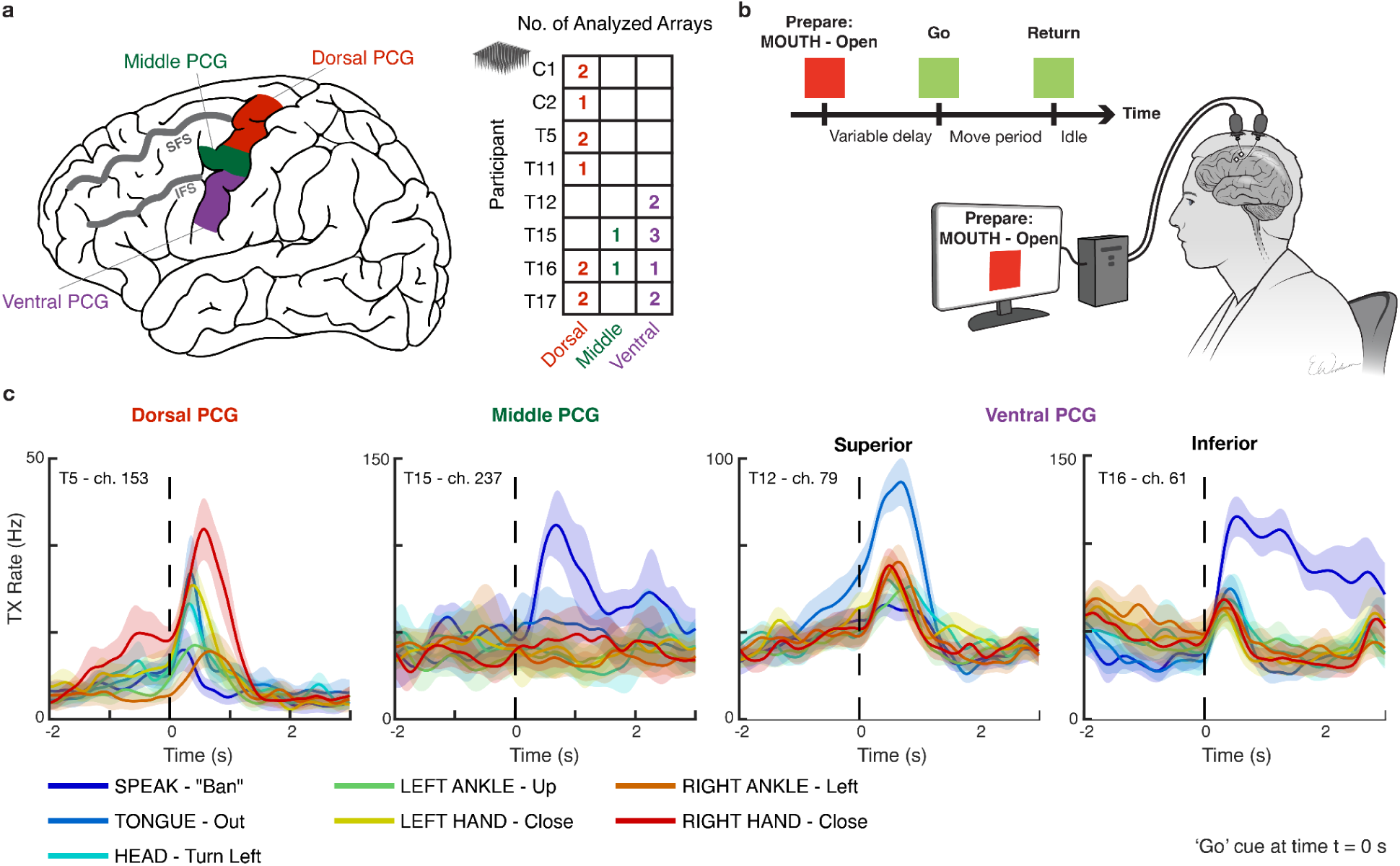
Sampling the length of precentral gyrus to assess whole-body movement representation. **a,** Diagram of the dorsal, middle, and ventral regions of precentral gyrus (PCG) sampled in this study. Neural activity was analyzed from a total of 20 microelectrode arrays distributed across 8 participants (right table). Four additional microelectrode arrays were recorded but excluded from analysis due to lack of tuning to any movement (see Methods). All arrays were placed in left precentral gyrus (and all participants except T12 and T15 were right-handed). **b,** Neural tuning to 46 attempted movements across the body was evaluated for each participant using an instructed delay task. **c,** Example responses from microelectrodes within the three sampled regions of PCG which show broad tuning to speech and movements of the arms, legs, face, and head. Each line shows the mean threshold crossing (TX) rate across all trials of a single movement condition and shaded regions show 95% confidence intervals (CIs). TX rates are a measure of spiking activity on an electrode and were denoised by convolving with a Gaussian smoothing kernel (120 ms s.d.).

### Neural tuning to the whole body in PCG

We assessed neural tuning to speech and movements of the face, head, arms and legs in eight participants in a visually cued movement task (Fig. 1b). If participants could not physically complete a movement due to paralysis, they were instructed to attempt to complete the movement as best as possible. Example electrodes demonstrating strong neural tuning to a variety of movements are shown in Fig 1c.

To quantify neural tuning at a coarse scale, we measured the degree to which the neural population activity evoked by each movement differed from a ‘Do Nothing’ baseline condition (Fig. 2a). The majority of electrode arrays (15 of 20) recorded robust neural modulation in response to *all* tested movements, with the remainder showing modulation to almost all movements. We also analyzed neural tuning at an individual electrode level and found that intermixed tuning to all body regions was frequently present within single electrodes as well (Extended Data Fig. 1).

**Fig. 2.**
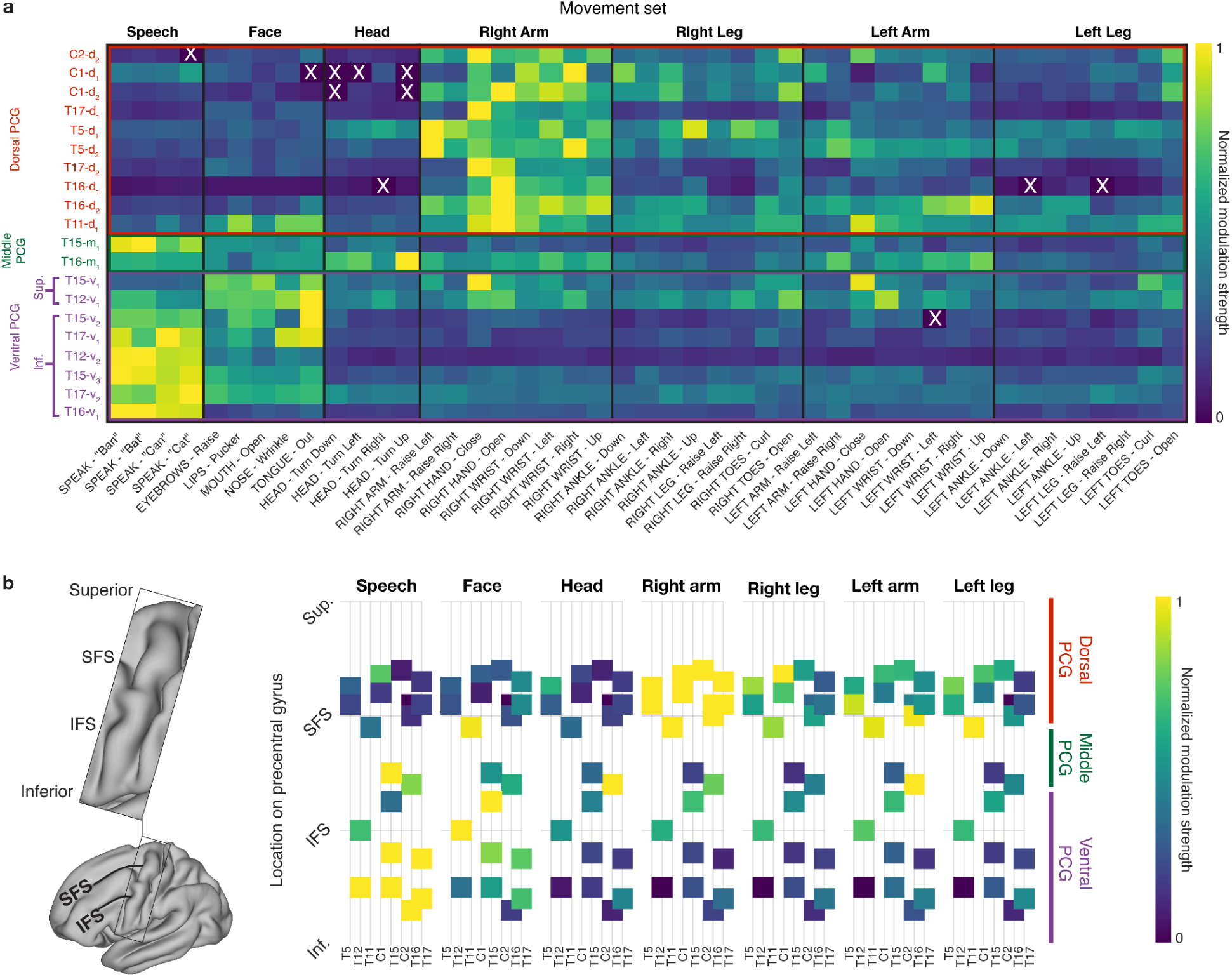
Regional organization of whole-body movement tuning in PCG. **a,** Neural modulation strength for each attempted movement was quantified by comparison to a ‘Do Nothing’ condition within each electrode array. The matrix is row-wise normalized to better visualize relative modulation strength (brighter colors indicate larger differences between movement-related activity and rest activity). A white ‘X’ denotes non-significant modulation. In general, each array features broad tuning to all movements, with region-specific trends in tuning strength. Arrays are coded by participant identifier and region (d=dorsal, m=middle, v=ventral), with subscripts that order multiple arrays from the same participant in a medial-to-lateral sequence. All arrays were placed in the left hemisphere. **b,** Approximate spatial organization of arrays along PCG relative to the anatomical landmarks of the superior and inferior frontal sulci (SFS and IFS, respectively). Each array (square) is colored in relation to its normalized modulation strength for a given movement class. Arm-preference is a general feature near the SFS (canonical hand-knob area), while speech-preference is seen near the inferior and middle regions of PCG, flanking a more broadly tuned orofacial region near the IFS (canonical face/head area).

Although all regions of PCG were tuned to the whole body, sorting the arrays by anatomical location revealed larger neuronal firing rate modulation aligned with the canonical motor homunculus, with arrays in dorsal PCG showing a preference for arm movements, and arrays in ventral PCG preferring orofacial movements (Fig. 2b). However, two findings also diverged from the classical homunculus. First, a majority of the arrays in the canonical orofacial region preferred speech to orofacial movement, particularly the most inferior ventral arrays (e.g., T16-v_1_) and one middle PCG array (T15-m_1_). Second, the 3 arrays in between these speech-preferring arrays were more broadly tuned to the whole body. Arrays T12-v_1_ and T15-v_1_ in particular had substantial limb tuning and were also most strongly activated by orofacial movements as opposed to speech.

### Neural decoding of the whole body from single sites in PCG

Next, to investigate the extent to which the whole body is represented in a *movement-specific* way throughout PCG, we tested whether a recurrent neural network decoder could classify between movements belonging to the same category using neural activity from single trials (Fig. 3). With the exception of speech, where four similar words could not be classified above chance in dorsal arrays, all categories of movement could be classified above chance in all arrays (Fig 3a). This indicates that a *differentiable* whole-body representation exists at each point in human motor cortex, but with a different emphasis depending on the location (with dorsal PCGshowing greater decodability of limb movements, and ventral PCG showing greater orofacial and speech decodability).

**Fig. 3.**
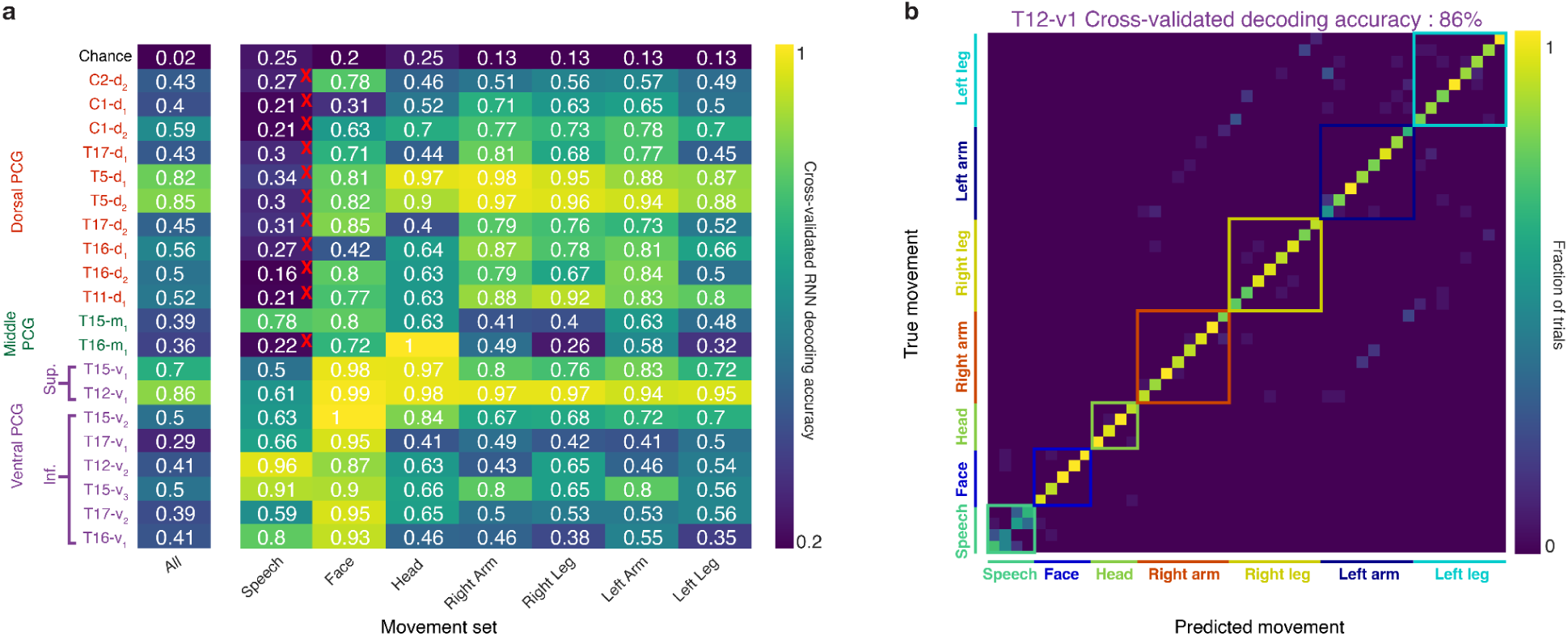
PCG contains a decodable whole-body representation at each sampled location. **a,** Heatmap depicting classification accuracies from a recurrent neural network decoder trained to classify movements using single trial neural recordings from each array separately. Red “X” denotes classification accuracies not significantly above chance. With the exception of speech in dorsal arrays, *all* movement categories can be decoded above chance in *all* arrays. Of note, superior-ventral arrays T15-v_1_ and T12-v_1_ express some of the highest decoding accuracies when classifying across all 45 movements. **b,** Example confusion matrix for superior-ventral array T12-v_1_, showing high accuracy across all movement types despite being a canonically orofacial area.

Notably, the two most superior ventral arrays (T12-v_1_, T15-v_1_) had some of the highest classification accuracies across all movement types. Fig. 3b shows an example confusion matrix between all 45 movements using activity from array T12-v_1_, which had an 86% decoding accuracy. This result further supports the idea of a superior-ventral orofacial zone with broad tuning to the whole body, but relatively weaker tuning to speech (as compared to the speech zones in middle PCG and inferior ventral PCG).

When interpreting these classification results, it is important to keep in mind two potential confounds: (1) some arrays simply record more tuned neurons than others, increasing their classification accuracy in general, and (2) some categories may be harder to classify than others due to the particular selection of movement conditions (e.g. speech, where the 4 tested words are very similar to each other). The second confound could explain why classification accuracy for speech is generally lower than other categories of movement, in contrast to the normalized firing rate modulation results above, where many arrays were speech-dominant.

### Four functional zones in PCG

Next, we applied principal components analysis (PCA) to the normalized firing rate modulation profiles and classification accuracies of each array to visualize array tuning characteristics in a low-dimensional space (Fig. 4a). PCA identified two major axes that explained 71% of the variance in tuning profiles: a “speech/face vs. limb” axis that discriminated the inferior ventral arrays (and middle PCG array T15-m_1_) from the dorsal arrays, and a “breadth of tuning” axis that discriminated the superior-ventral arrays from the others. The coefficients for these two PCA axes (Fig. 4b) were similar for both the firing rate modulation profiles (solid lines) and classification accuracies (dashed lines), indicating general agreement between these two complementary methods of assessing whole-body tuning.

**Fig. 4.**
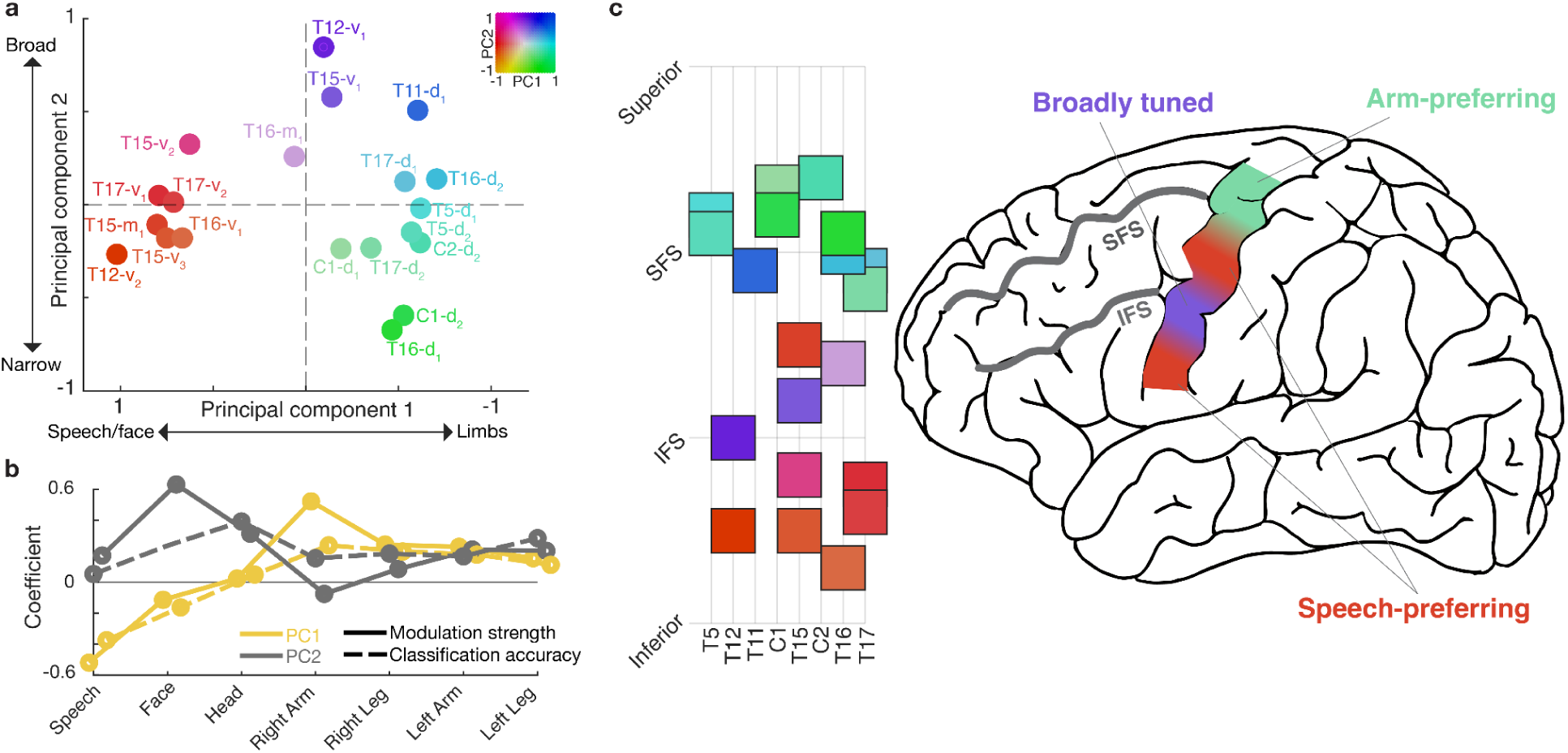
Four functional zones in PCG. **a,** Low-dimensional representation of the tuning properties of each array, generated by applying principal components analysis to the neural tuning magnitudes and classification accuracies associated with each array. Arrays are colored according to their position in PCA space (colors are used to construct the topographical map of tuning in panel c). **b,** Coefficients of the top two principal components. The first principal component describes a speech/face vs. limb axis, while the second describes breadth of tuning. Solid lines denote coefficients related to modulation strength and dashed lines denote coefficients related to classification accuracies. **c**, Arrays are shown topographically and colored according to their position in PCA space (colors match **a**), revealing functional zones of tuning properties. Combining this information with what is known in the literature, we hypothesize that there are four functional zones in the sampled region of PCG: an inferior-ventral and middle zone related to speech (orange), a superior-ventral orofacial area that also contains strong arm/hand tuning (purple), and one arm/hand dominant zone (cyan).

We colored each array according to its location along the top two PCA axes, and displayed this information topographically to reveal regional patterns in tuning characteristics (Fig. 4c). Based on these results, as well as what is known from prior work (particularly studies identifying middle PCG as a speech/language region^15–17^), we hypothesize that there are four main functional zones in the sampled area of PCG (Fig. 4d): one dorsal zone emphasizing arm movements, one inferior-ventral zone emphasizing speech, one middle PCG zone also emphasizing speech (array T15-m_1_), and one superior-ventral orofacial zone between the speech zones that is most broadly tuned to the whole body (arrays T12-v_1_ and T15-v_1_). We also analyzed fMRI resting state data from the Human Connectome Project^16^ and found evidence of two resting state networks that parcellate PCG into four zones (Extended Data Fig. 2): one language-related resting state network with hotspots in inferior-ventral PCG and middle PCG (accounting for the speech zones), and one hand/arm-related resting state network with hotspots in dorsal PCG and superior-ventral PCG (which could explain the strong limb tuning in superior-ventral PCG).

It is possible that anatomical variability across individuals could cause other regions to sometimes appear on PCG. Array T16-m_1_ in particular appears to be an outlier in exhibiting strong head-related modulation and classification accuracy. Using the Human Connectome Project cortical parcellation procedure applied to T16’s pre-operative neuroimaging data, we identified array T16-m_1_ as being on the border of the premotor eye field, an area which may be highly tuned to movements of the eyes and head (Extended Data Fig. 3).

### A compositional whole-body neural code

Finally, we sought to elucidate the representational structure of the whole-body code found throughout PCG. In prior work investigating hand knob area^11^, we found a “compositional” neural code linking all four limbs together that had two main features: (1) correlated representations of homologous movements across limbs (e.g., wrist flexion and ankle flexion), and (2) representation of the limb itself independent of the movement.

Here, we also found strong correlations between all four limbs in multiple regions of PCG. Fig. 5a shows example arm/leg correlations for dorsal PCG and superior-ventral PCG (Extended Data Fig. 4 shows correlation matrices for all regions and limb pairs). Representational similarity between limbs was present for most limb pairs and most arrays (Fig. 5b), although the arms appeared to be most strongly correlated with one another (significant for 17 of 20 arrays). These results show that inter-limb neural correlations linking all four limbs are a general feature of human motor cortex, appearing even in orofacial and speech regions.

**Fig. 5.**
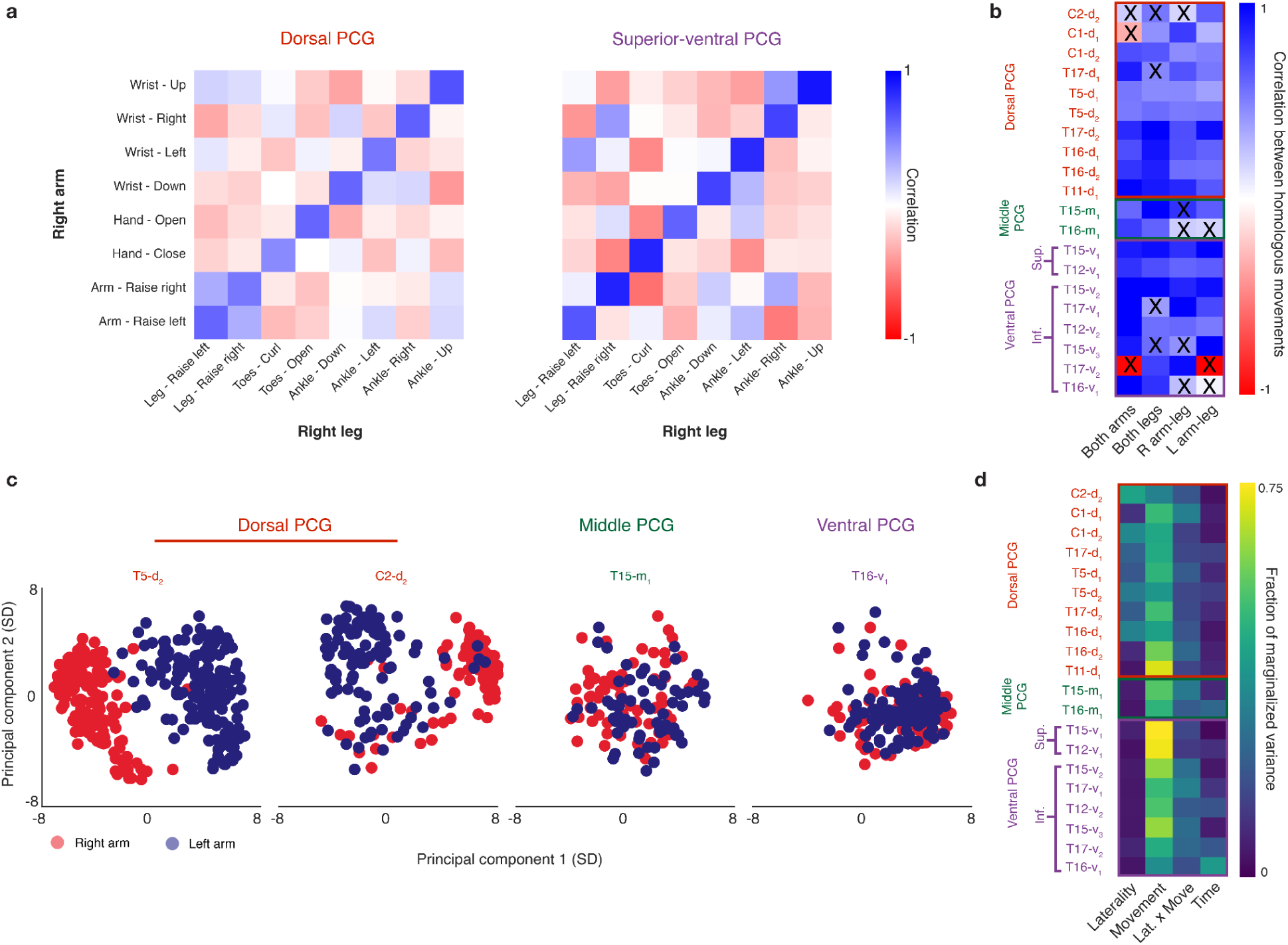
A compositional neural code throughout PCG that links all four limbs together. **a,** Representational similarity of neural activity for homologous arm and leg movements in two example regions of PCG. Each square in the matrix represents pairwise similarity for two movements (as measured by the correlation between neural firing rate vectors). Ordering movements according to their arm-leg homolog shows a diagonal structure indicating high similarity between homologous movements. **b,** Summary of representational similarity for each electrode array across various limb pairs. Each square in the matrix represents the average similarity between homologous movements of a limb pair (as measured by the mean of the diagonal values of the corresponding similarity matrix). A black ‘X’ indicates non-significant similarity via shuffle testing. **c,** Low-dimensional PCA visualizations of single-trial neural activity for 4 example arrays (each trial is a single circle that is colored according to the side of the body; only arm movement conditions are plotted for clarity). The dorsal arrays show a clear “laterality dimension” that separates the right and left arms, whereas the middle and ventral arrays feature a more intermixed representation in the top 2 PCs. **d,** Summary of the size of the laterality-related variance for arm movements relative to other factors of the neural data. Data was marginalized according to the following factors: time, laterality, movement, and the laterality-movement interaction. Generally, the laterality dimension was stronger for the dorsal arrays in comparison to the middle and ventral arrays (as measured by fraction of variance within the marginalization).

Next, we tested for the existence of limb-coding activity in the form of ‘laterality’ neural dimensions that would separate the left from right arms independently of the movement itself, as found previously in hand knob area^11,18^. Consistent with prior results, we found that a laterality dimension is a dominant feature of dorsal PCG, appearing clearly in the top two principal components of the neural activity (Fig. 5c; Extended Data Fig. 5 shows scatterplots for all arrays). Laterality information was less prevalent in other areas of PCG (Fig. 5c,d), indicating that this feature is likely unique to dorsal PCG.

## Discussion

Penfield’s motor homunculus has influenced our understanding of how the brain controls movement and has shaped decades of motor neuroscience studies. Penfield and Rasmussen noted that the homunculus was intended as "an aid to memory" and not a precise parcellation of cortex, stating "it is a cartoon of representation in which scientific accuracy is impossible." Despite this caveat, there still exists considerable uncertainty in the literature about the degree to which PCG in humans adheres to a homuncular somatotopy^3–6,11,13,14,19–22^. Over the past few years, recent studies using fMRI and ECoG^13,14^ have highlighted specific regions of intermixed tuning in motor cortex inconsistent with the homuncular model, but the extent to which intermixing occurs at a single neuron level was not known.

Here, a comprehensive motor map of PCG at single neuron resolution, leveraging microelectrode array recordings from 20 arrays across 8 individuals, revealed that differentiable, whole-body representations exist at *all* sampled points of PCG. This indicates that motor representations are substantially more diffuse and intermixed than some previously reported exceptions have shown^20,23–26^. We also found that these whole-body representations are abstract, linking together the legs and arms in a partially limb-independent representation of motor action. A limb-independent movement code could facilitate the transfer of learned actions from one limb to another^27–33^, and was present throughout all sampled areas of PCG. The consistency of results shown across a wide range of disorders (spinal cord injury, ALS and brainstem stroke) and degree of motor impairment suggest that these results reflect a general property of human motor cortex as opposed to reorganization due to disease, although only recordings from able-bodied individuals can provide a definitive answer.

Our results also shed new light on the regional arrangement of human motor cortex. We found evidence that the sampled extent of PCG between the sylvian fissure and superior frontal sulcus is composed of four functional zones, each with their own whole-body representations: (1) a dorsal arm/hand zone, (2) a middle speech zone (also highlighted in recent work^15–17^), (3) an inferior-ventral speech zone, and (4) a broadly-tuned, superior-ventral orofacial zone. The superior-ventral region was strongly tuned to both orofacial and arm/hand movement, and may be homologous to macaque ventral premotor cortex, which is known to contain grasping, reaching and orofacial related information^34–36^ (see also^37^). The middle and inferior-ventral speech zones have previously been interpreted as motor regions that control the larynx^38,39^. However, these regions are correlated with known language areas in resting state fMRI data (Extended Data Fig. 2), and in our prior work, offered much higher speech decoding performance relative to the superior-ventral region^40,41^ (even for unvoiced phonemes^41^). These areas may thus be related to speech more generally, as opposed to being involved only in aspects of speech related to laryngeal control.

Although motor representations in the human brain have been an active area of investigation for decades^3,5,11,13,14,23,26,36,42–48^, our findings here suggest that low-resolution imaging modalities and electrical stimulation have historically overemphasized the extent to which the body is segregated in motor cortex. Nevertheless, it is still possible that more classically homuncular body representations exist deep within the central sulcus. Anatomical studies identify Brodmann area 4 (primary motor cortex) as typically lying within the central sulcus, while PCG is thought to be comprised largely of Brodmann area 6 (premotor cortex)^49–53^. In macaques, primary motor cortex is thought to contain more segregated^54^ and less abstract movement representations^55,56^. A recent cortical atlas based on neuroimaging data from the Human Connectome Project^16^ also divides PCG into a series of premotor areas that are distinct from the central sulcus. Visualizing task activation fMRI from this atlas reveals a sharper separation of foot, hand and tongue in central sulcus that is more mixed in PCG (Extended Data Fig. 6). Future studies obtaining single neuron recordings from the central sulcus can provide a definitive answer.

Finally, these results chart the first map of how well each movement category can be decoded at single neuron resolution throughout PCG. This map can inform the design of intracortical brain-computer interfaces (BCIs) aiming to restore speech, arm, hand and leg movement to people with paralysis^40,41,57–60^. To achieve the highest performance with the minimal number of electrodes, our results show that speech BCIs may benefit from targeting inferior-ventral and middle PCG while avoiding the superior-ventral region. In contrast, the superior-ventral region may be an underappreciated complement to dorsal PCG for arm/hand BCIs. Overall, the widespread decodability of the whole body throughout PCG is advantageous for BCIs seeking to restore multiple functions with a small footprint.

## Data Availability

All data required to reproduce these results will be publicly released upon acceptance in a peer-reviewed journal.

## Code Availability

Code for cross-validated estimation of neural correlation and distance is available at: https://github.com/fwillett/cvVectorStats. Code for implementing the recurrent neural network decoder is available at: https://github.com/cffan/neural_seq_decoder.

## Acknowledgements

We thank participants T5, T11, T12, T15, T16, T17, C1, and C2 and their caregivers for their generously volunteered time and dedicated contributions to this research; B. Davis, K. Tsou and S. Kosasih for administrative support; and Angelique Paulk and Peter Hadar for hep with T17 presurgical imaging. Support provided by the NIH National Institute of Neurological Disorders and Stroke (U01NS123101); NIH National Institute on Deafness and Other Communication Disorders (R01DC014034); Wu Tsai Neurosciences Institute; Howard Hughes Medical Institute; Larry and Pamela Garlick; Office of Research and Development, Rehabilitation R&D Service, US Department of Veterans Affairs (A2295R, N2864C); ; NIH National Institute on Deafness and Other Communication Disorders (K23DC021297); AHA’s Second Century Early Faculty Independence Award;

* The contents do not represent the views of the Department of Veterans Affairs or the US Government.

CAUTION: Investigational Device. Limited by Federal Law to Investigational Use.

## Author Information

These authors jointly supervised this work: Jaimie M. Henderson and Francis R. Willett

## Author Contributions

DRD and FRW conceived the study, wrote the manuscript, and led the development, analysis, and interpretation of all experiments.

EYC and DRD applied the Human Connectome Project cortical parcellation procedure for participants T12, T15, T16, and T17. DQ and YM helped facilitate and develop scanning protocols for T16.

EVO, ALP, SRN, TH, NSC, MW, and JJ developed and executed the experiments at their respective sites.

NH, FBK, CN, CI, SRN, and AA were responsible for coordination of session scheduling, logistics and daily equipment setup/disconnection for participants T5, T12, T11, T15, T16, and T17 respectively.

LRH is the sponsor-investigator of the multisite BrainGate2 pilot clinical trial.

DBR, LRH, SSC and ZMW planned T17’s array placement surgery, and ZMW performed T17’s array placement surgery. DBR and LRH were responsible for all clinical trial related activity at MGH. DBR and LRH supervised and guided all research activity with T17.

LRH, SSC and ZMW planned T11’s array placement surgery, and ZMW performed T11’s array placement surgery. LRH was responsible for all clinical trial related activity at VA Providence and MGH. LRH supervised and guided all research activity with T11.

JMH planned and performed T5’s and T12’s array placement surgery and was responsible for all clinical trial-related activities at Stanford.

DMB planned and performed T15’s array placement surgery and was responsible for all clinical trial-related activities at UC Davis. SDS and DMB supervised and guided all research activities at UC Davis.

NAY planned and performed T16’s array placement surgery and was responsible for all clinical trial-related activities at Emory. CP and NAY supervised and guided all research activities as Emory.

JED and SJB planned C1 and C2’s array placements and were responsible for all clinical trial-related activities at the University of Chicago.

The study was supervised and guided by JMH and FRW.

All authors reviewed and edited the manuscript.

---

**Extended Data Fig. 1.**
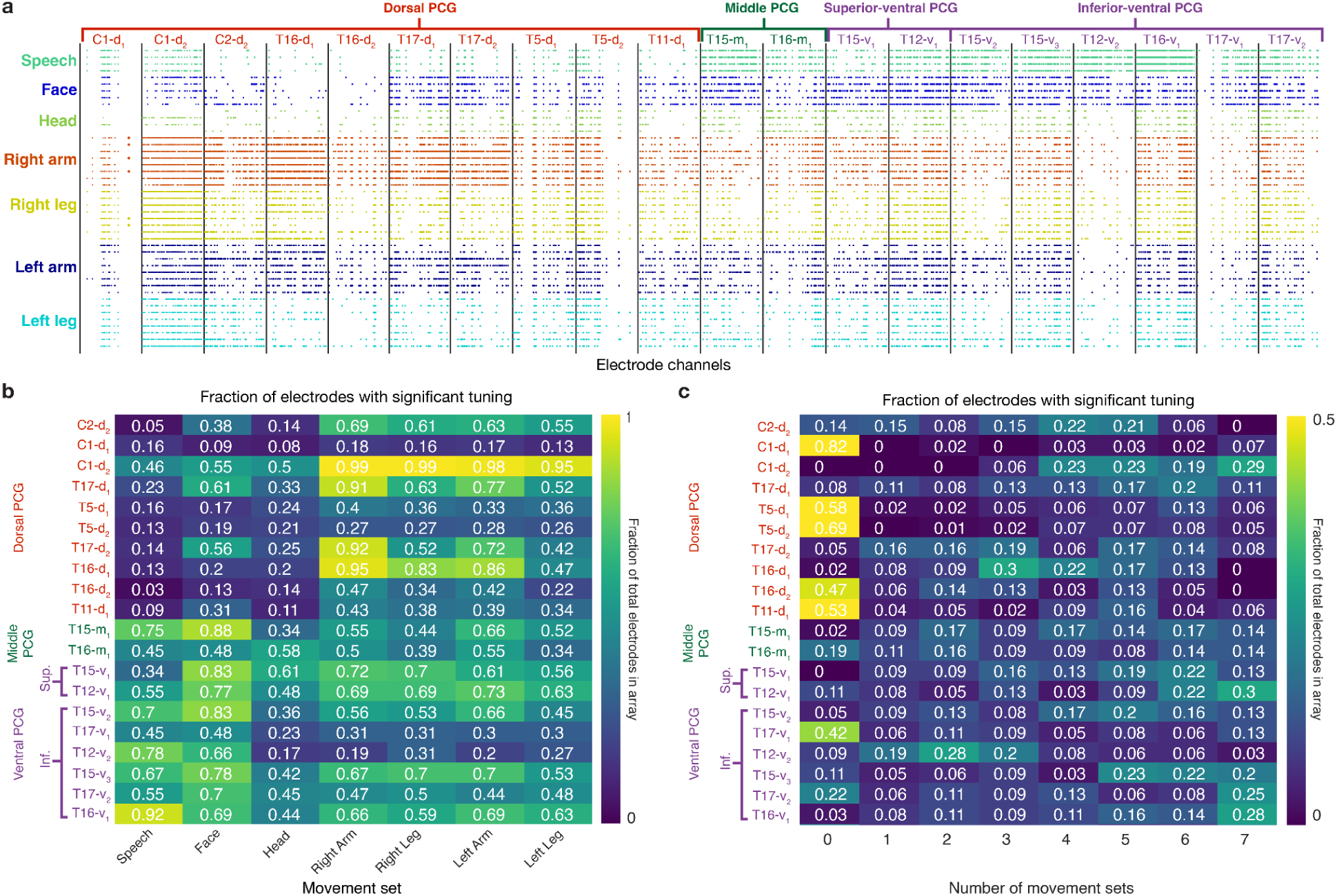
Tuning to speech, face, head, arm, and leg movements is intermixed within single electrodes. **a,** A tuning significance matrix that indicates whether an electrode was significantly tuned to each individual movement (rows). A colored circle (i,j) within the matrix indicates significant tuning of electrode j to movement i. The circle color corresponds to the associated movement category. Significant tuning was assessed by comparing the firing rates for each movement against the ‘Do Nothing’ condition using a 2-sample t-test. P values < 0.00001 were considered significant. All electrode arrays contain electrodes with intermixed tuning to each movement set. **b,** Heat map summarizing the fraction of electrodes within each array that were significantly tuned to each movement category as computed in **a**. If an electrode showed significant tuning to at least one movement within each movement set, it was considered significantly tuned to that movement set. **c,** Heat map summarizing the fraction of electrodes that had statistically significant tuning to each possible number of movement sets (from 0 to 7). Many electrodes are tuned to more than one movement set, indicating intermixed tuning to the whole-body at a single electrode level.

**Extended Data Fig. 2.**
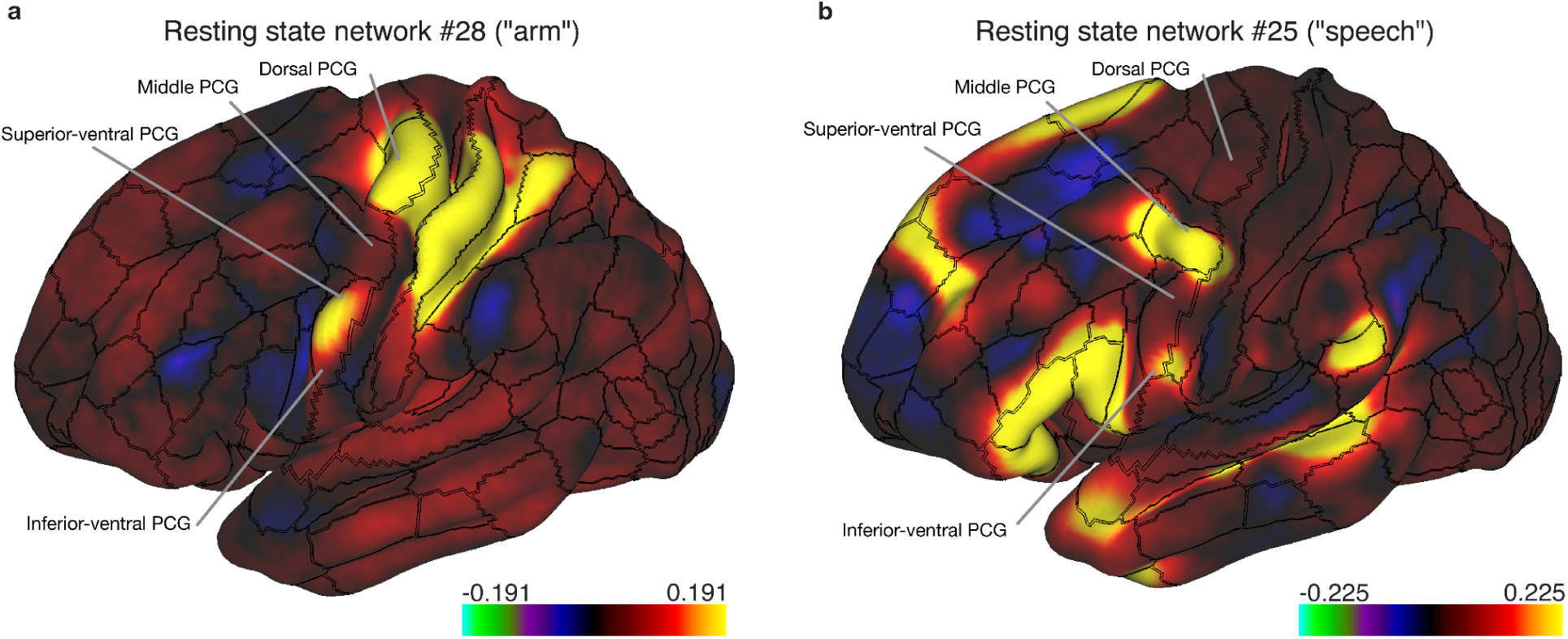
Resting state fMRI networks identified with a large-sample size dataset (n=210) support the idea of four functional zones in PCG. Group-averaged resting state networks from the Human Connectome Project (HCP) identify sets of cortical areas with correlated activity during resting state. **a,** “Network 28” shows that superior-ventral PCG appears connected to dorsal PCG (hand knob) and areas of somatosensory cortex known to be related to the hand/arm, supporting the idea of superior-ventral PCG being a distinct functional zone with strong hand/arm tuning in addition to orofacial tuning. **b,** “Network 25” shows that middle PCG and inferior-ventral PCG appear to be connected to a larger/speech language network, including Broca’s area and Wernicke’s area, supporting the idea that these zones in PCG are speech-specific.

**Extended Data Fig. 3.**
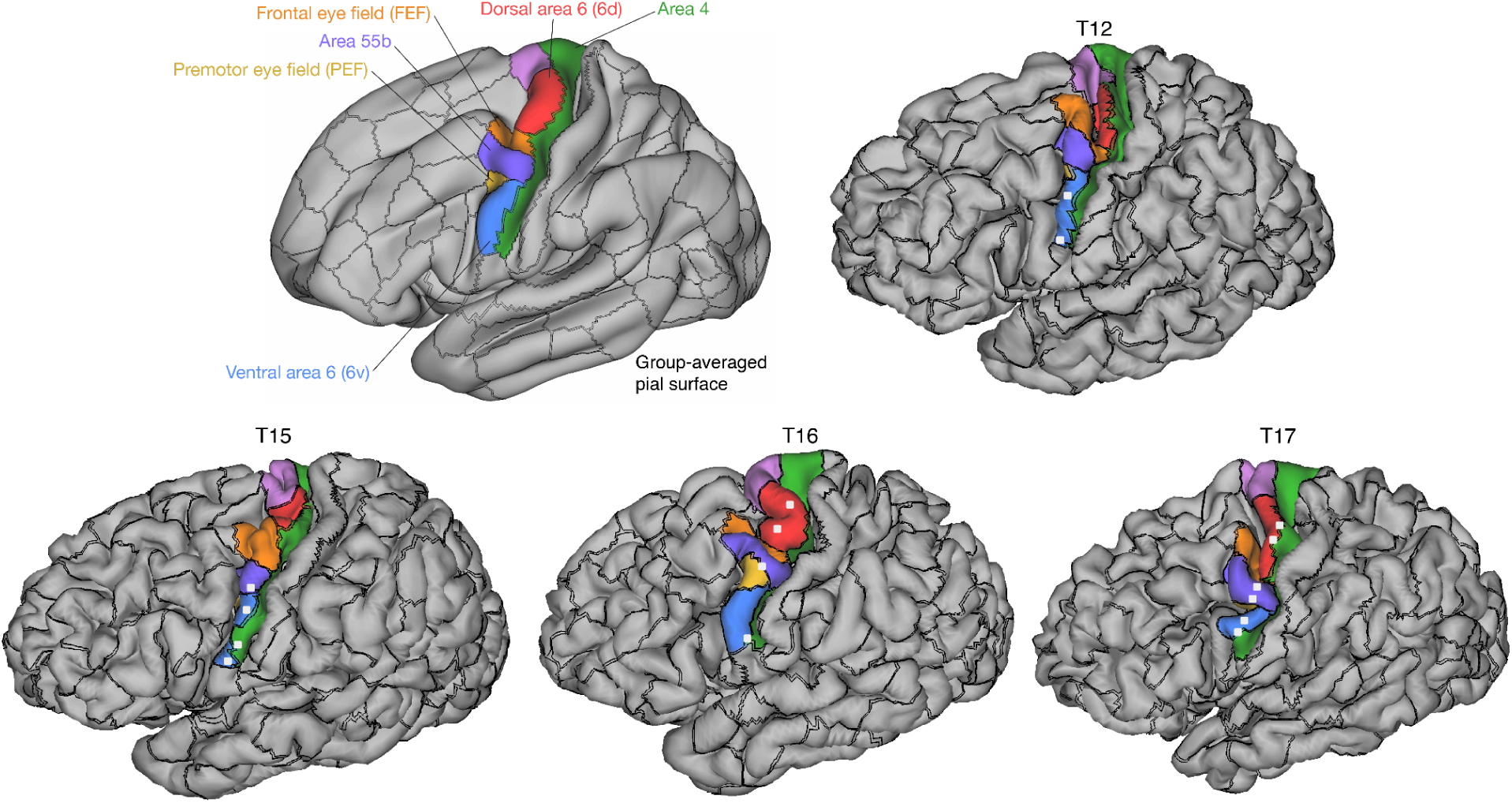
Human Connectome Project (HCP) cortical parcellations for participants where HCP-style imaging data was collected prior to array implantation. Cortical parcellations for participants T12, T15, T16, and T17 were performed using the HCP multi-modal parcellation pipeline. Each participant’s pial surface is pictured with areal borders (black lines) of the parcellated regions. Regions of interest along PCG are shown in color. White squares indicate array placement locations estimated using photographs taken during surgery.

**Extended Data Fig. 4.**
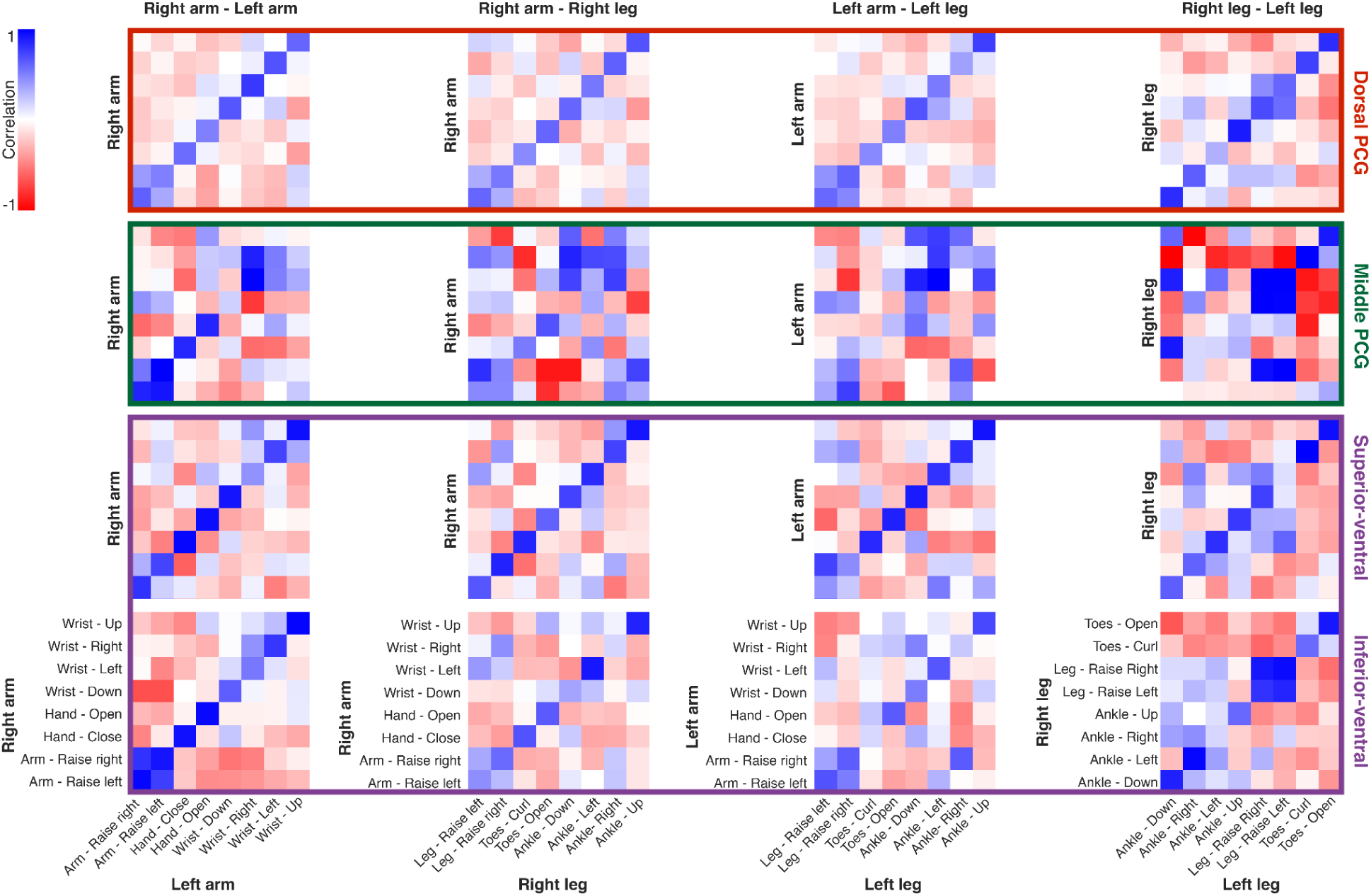
Representational similarity of homologous movements across limb pairs (columns) and brain regions (rows). Each square in a similarity matrix represents pairwise similarity for two movements (as measured by the correlation between neural firing rate vectors). Each matrix is the average similarity matrix across all arrays within that brain region (dPCG, mPCG, superior vPCG, or inferior vPCG), as indicated by the colored boxes. Movements are ordered such that the diagonal of each similarity matrix indicates the correlations between homologous movements. High similarity between homologous movements (dark blue diagonal bands) seems apparent in all regions except mPCG.

**Extended Data Fig. 5.**
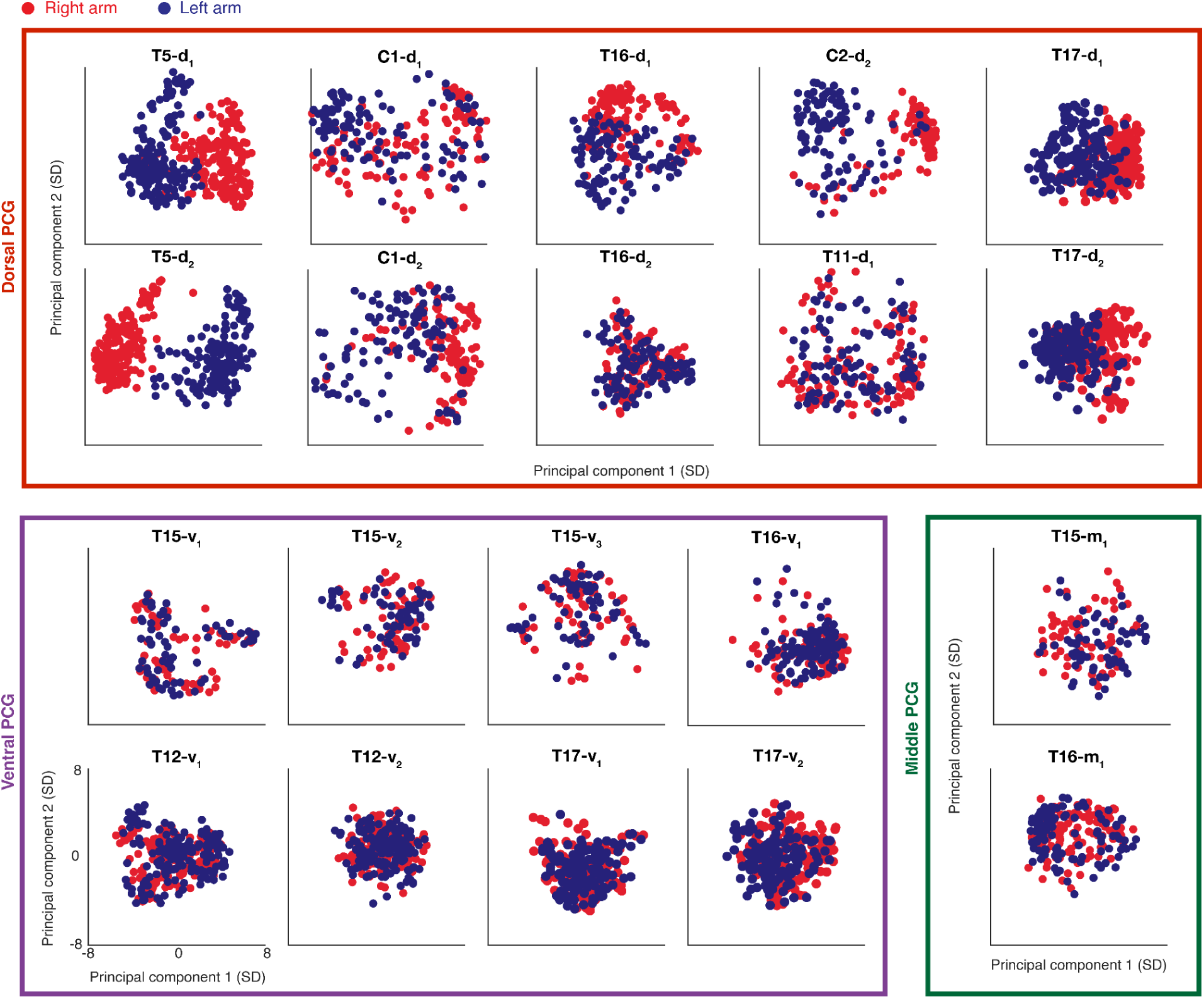
A laterality-related neural dimension is a dominant feature of dorsal PCG. Same as Figure 3c, with the low-dimensional PCA visualizations of single-trial neural activity plotted for each array. Most dorsal arrays show a clear “laterality dimension” that separates the right and left arms, whereas all arrays in middle and ventral PCG feature a more intermixed representation in the top 2 PCs.

**Extended Data Fig. 6.**
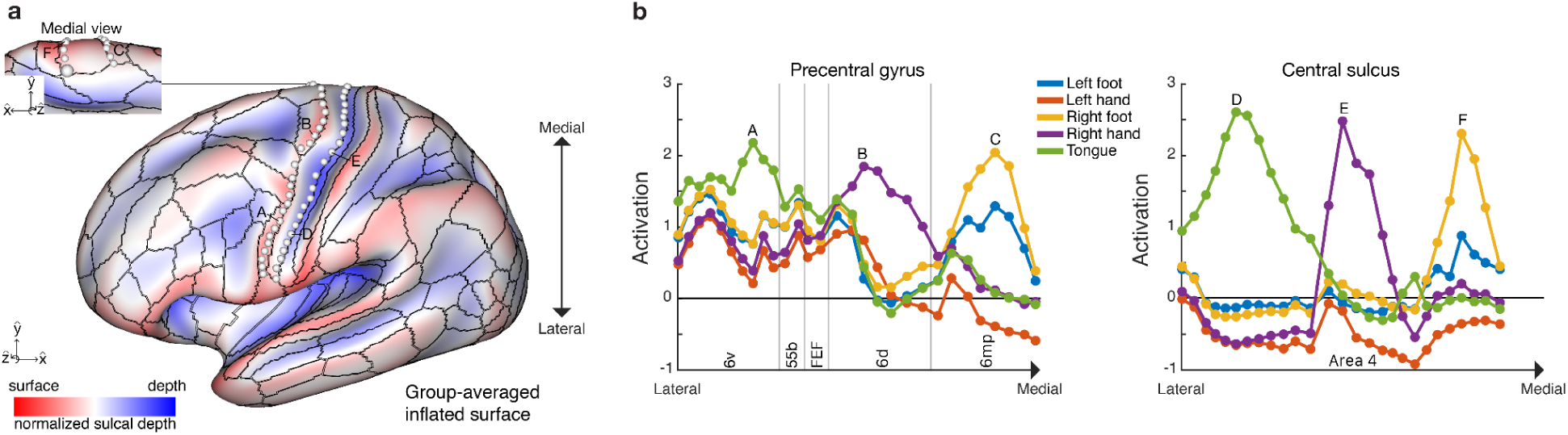
Task fMRI activation data from the Human Connectome Project (HCP) large-sample dataset (n=1,200) supports the idea of more segregated body representations in the central sulcus. **a,** HCP group-averaged inflated brain surface with a colored gradient indicating sulcal depth (surface as red, depth as blue). White circles along the fundus of the central sulcus and crest of PCG indicate probed areas for motor activation from the group-averaged task-fMRI. **b**, Task-fMRI activations for the probed areas in **a.** The depths of the central sulcus feature more selective activation for individual body parts in comparison to the crest of PCG which shows more intermixed tuning.

**Extended Data Fig. 7.**
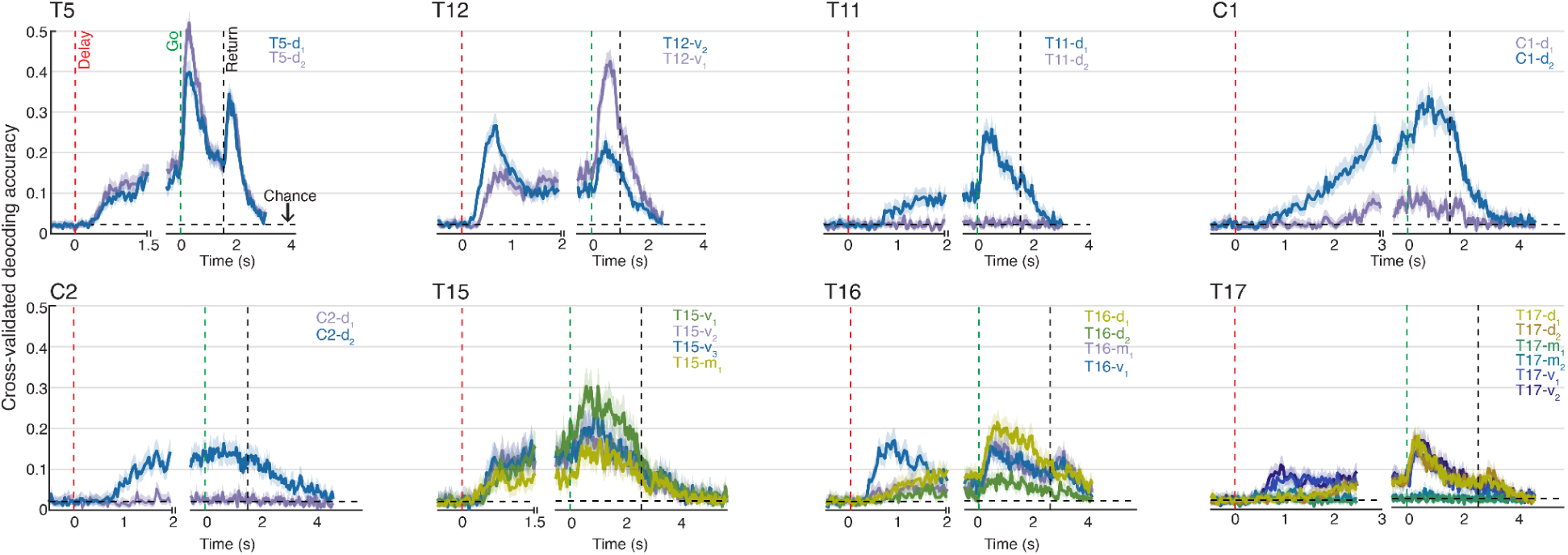
Neural decoding accuracy across all movements using a sliding window (used as a criterion for array exclusion). Movement classification sweeps for all arrays, which was used as a criterion to exclude arrays from analysis if classification accuracy was never consistently above chance. Lines represent classification accuracy (across all 46 movement conditions) across time for each participant’s arrays. Classification was performed with a cross-validated Gaussian naive Bayes classifier applied to a sliding window of 100 ms of neural population activity. Due to the variation in delay periods, classification was performed in two sets, the first aligned to the cue onset (‘Delay’ cue) and the other aligned to the ‘Go’ cue (resulting in a discontinuity in the time axis). Shaded regions show 95% CIs. Vertical dashed lines indicate the onsets for each cue (‘Delay’ cue in red, ‘Go’ cue in green, and ‘Return’ cue in black). Chance level decoding performance (1/46) is indicated by the horizontal dashed black line. Arrays with non-significant decoding performance (CIs intersected chance level throughout all time periods) were excluded from this study (arrays T11-d_2_, C2-d_1_, T17-m_1_ and T17-m_2_).

## Supplementary Methods

## 1 Experimental procedures

### 1.1 Study participants

This study includes data from eight participants who each gave informed consent prior to any experimental procedures. Participants T5, T11, T12, T15, T16, and T17 were enrolled in the BrainGate2 Neural Interface System clinical trial (ClincialTrials.gov Identifier: NCT00912041, registered June 3, 2009). This pilot clinical trial was approved under an Investigational Device Exemption (IDE) by the US Food and Drug Administration (Investigational Device Exemption #G090003). Permission was also granted by the Stanford University Institutional Review Board (IRB; protocol #20804), the Mass General Brigham IRB (protocol #2009P000505), the University of California, Davis IRB (protocol #1843264), the Emory University IRB (protocol #00003070), and Providence VA Healthcare IRB. Participants C1 and C2 were enrolled under a separate multi-site clinical trial (ClincialTrials.gov Identifier: NCT01894802, registered July 10, 2013) which was also conducted under an IDE from the US Food and Drug Administration and approved by the IRBs at the University of Pittsburgh and the University of Chicago. All research was performed in accordance with relevant guidelines and regulations.

T5 is a right-handed man, 69 years of age at the time of this study, with tetraplegia due to cervical spinal cord injury (classified as C4 AIS-C) which occurred approximately 9 years prior to enrollment in the clinical trial. T5 has two 96-channel intracortical microelectrode arrays (Blackrock Microsystems, Salt Lake City, UT; 1.5 mm electrode length) placed in the hand knob area of the left precentral gyrus (PCG). The hand knob area was identified anatomically by preoperative magnetic resonance imaging (MRI). T5 has full movement of the face and head and the ability to shrug his shoulders. Below the level of spinal cord injury, T5 has very limited voluntary motion of the legs and arms.

T11 is a right-handed man, 38 years of age at the time of this study, with tetraplegia due to a cervical spinal cord injury (classified as C4 AIS-B) which occurred approximately 14 years prior. T11 has two 96-channel intracortical microelectrode arrays (Blackrock Microsystems; 1.5 mm electrode length) placed in the left dorsal PCG, targeting the hand knob area as identified anatomically by preoperative MRI. T11 has full movement of the face and head with very limited voluntary motion of the arms.

T12 is a left-handed woman, 67 years of age at the time of this study, with slowly-progressive bulbar-onset Amyotrophic Lateral Sclerosis (ALS). T12 was diagnosed at age 59 with an ALS-FRS score of 26 at the time of study enrollment. T12 has four 64-channel intracortical microelectrode arrays (Blackrock Microsystems; 1.5 mm electrode length) placed in the left (language dominant, motor non-dominant) hemisphere, based on preoperative anatomical MRI, functional MRI and cortical parcellation using the Human Connectome Project^16^ (HCP) multi-modal parcellation pipeline (see Extended Data Figure 6 for cortical parcellation results).

Two arrays were placed in the inferior frontal gyrus (not included in this study), and two arrays were placed in the ventral PCG, targeting area 6v. See Willett et al. for more details^40^. T12 remains functionally independent with 3-4/5 strength in all limbs, but is anarthric (able to vocalize, but unable to produce intelligible speech).

T15 is a left-handed man, 45 years old at the time of this study, with ALS (ALS-FRS score of 23 at the time of study enrollment). T15 has four 64-channel intracortical microelectrode arrays (Blackrock Microsystems; 1.5 mm electrode length) placed in the left (language dominant, motor non-dominant) PCG, based on preoperative anatomical MRI and HCP cortical parcellation (see Extended Data Figure 6 for cortical parcellation results). One array was placed targeting area 55b, two arrays targeting area 6v, and one array targeting area 4. See Card et al. for more details^61^. T15 has limited orofacial movement with the capacity for vocalization, but is unable to produce intelligible speech. T15 has very limited voluntary motion of the rest of the body.

T16 is a right-handed woman, 52 years of age at the time of this study, with tetraplegia and dysarthria due to a pontine stroke approximately 19 years prior to enrollment in the BrainGate2 clinical trial. T16 has four 64-channel intracortical microelectrode arrays (Blackrock Microsystems; 1.5 mm electrode length) placed in her left precentral gyrus: two in the hand knob area (targeting area 6d), one in speech-related ventral premotor cortex (targeting ventral 6v), and one in middle precentral gyrus (targeting area 55b). Implant area targets were identified by the HCP multimodal cortical parcellation procedure (see Extended Data Figure 6 for cortical parcellation results). Examination of post-implant array locations indicated that the middle precentral gyrus array appears to be on the border between area PEF and 55b. T16 is able to speak slowly and quietly, but speech cadence is reduced due to poor diaphragm voluntary control. She has limited voluntary control of her upper extremities, with some shoulder motion and some slow and contractured wrist and finger movements. She has limited to no voluntary control of her lower extremities. T16’s sensation is fully intact.

T17 is a right-handed, 33-year-old man with a history of rapidly-progressive ALS. T17 has six 64-channel intracortical microelectrode arrays (Blackrock Microsystems; 1.5 mm electrode length) placed in the left hemisphere, based on preoperative anatomical MRI, task-based functional MRI, and cortical parcellation using the Human Connectome Project (HCP) multi-modal parcellation pipeline. Two arrays were placed in the dorsal precentral gyrus (targeting area 6d), two arrays were placed in the ventral precentral gyrus (targeting area 6v), and two arrays were placed in area 55b. At the time of this study, T17 is quadriplegic, anarthric, and ventilator dependent. His only remaining volitional motor control is over his extra-ocular movements.

C1 is a right-handed man, 57 years old at the time of implant, who presented with a C4-level ASIA D spinal cord injury that occurred 35 years prior to implant. C1 has four microelectrode arrays (Blackrock Microsystems; 1.5 mm electrode length) placed in the left hemisphere. Two 96-channel microelectrode arrays were implanted in the arm and hand area of the motor cortex and two other 32-channel arrays (not included in this study) were implanted in the somatosensory cortex. Targeted array placement was based on functional neuroimaging (fMRI) of the participant attempting to make movements of the hand and arm, within the constraints of anatomical features such as blood vessels and cortical topography. See Greenspon et al. for more details^62^. C1 had no control of the intrinsic or extrinsic muscles of the right hand but retained the ability to move his arm with noted weakness in many upper limb muscles. He retained impaired, but largely functional, movement of the other limbs, full control of head and face movement, and could speak fluently. The data included here were collected 3.25 years post-implant.

C2 is a right-handed man, 60 years old at the time of implant, who presented with a C4-level ASIA D spinal cord injury and right brachial plexus injury that occurred 4 years prior to implant. C2 has four microelectrode arrays (Blackrock Microsystems; 1.5 mm electrode length) placed in the left hemisphere. Two 96-channel microelectrode arrays were implanted in the arm and hand area of the motor cortex and two other 32-channel arrays (not included in this study) were implanted in the somatosensory cortex. Targeted array placement was based on functional neuroimaging (fMRI and MEG) of the participant attempting to make movements of the hand and arm, within the constraints of anatomical features such as blood vessels and cortical topography^62^. C2 retained full control of his entire body except for right hand and arm movement. He could speak fluently. The data included here were collected 0.75 years post-implant.

### 1.2 Neural signal processing

For each participant, neural signals were recorded from the microelectrode arrays using the NeuroPort^TM^ system (Blackrock Microsystems). The signals were then analog filtered (4th order Butterworth with corners at 0.3 Hz to 7.5 kHz) and digitized at 30 kHz (250 nV resolution). The subsequent digital filtering and neural feature extraction methods differed between participants due to variations in the systems at different sites.

For T5, T11, and T12 the signals were decimated to 15 kHz and bandpass filtered between 250 Hz to 4900 Hz using a 4th order zero-phase non-causal Butterworth filter. Linear regression referencing (LRR) was then applied to further reduce ambient noise artifacts^63^ prior to spike detection. Spike threshold crossing detection was implemented using a -4.5 x RMS threshold applied to each electrode, where RMS is the electrode-specific root mean square of the time series voltage recorded on that electrode.

For T15, the signals were band-pass filtered between 250 Hz to 5 kHz using a 4th order zero-phase non-causal Butterworth filter and LRR was then used to reduce noise artifacts. Spike threshold crossing detection was implemented using a -4.5 x RMS threshold.

For T16, each electrode was high-pass filtered with a 250 Hz cutoff using a 4th order zero-phase non-causal Butterworth filter. LRR was used for noise reduction and artifact removal with parameters computed post-filtering from a dedicated reference block at the beginning of the session. A -3.5 x RMS threshold was applied to each electrode for spike threshold crossing detection.

For T17 the signals were decimated to 15 kHz and bandpass filtered between 250 Hz to 5000 Hz using a 4th order zero-phase non-causal Butterworth filter. Linear regression referencing (LRR) was then applied to further reduce ambient noise artifacts prior to spike detection. Spike threshold crossing detection was implemented using a -3.5 x RMS threshold applied to each electrode.

For participants C1 and C2, a high-pass filter (250 Hz) was applied to each electrode prior to spike detection. Spike threshold crossing detection was implemented using a -4.5 x RMS threshold applied to each electrode.

The resulting spiking data from each participant mentioned above was binned in 20 ms bins for offline analyses and decoding as presented throughout this study.

### 1.3 Overview of data collection sessions and cued movement task

For each participant, neural data was recorded in a single “session” on a scheduled day. During the session, the participant was seated in front of a computer monitor at an idle and relaxed position. Each participant completed a series of 5-10 minute “blocks” of the cued movement task, consisting of an uninterrupted series of trials. Table 1 lists all data collection sessions reported in this work. Variation in the number of trials and/or blocks collected for each participant is due to differences in session durations for each participant and their respective comfort/fatigue levels.

**Table 1.** List of data collection sessions.

| **Participant** | **RMS multiplier** | **Date** | **Blocks** | **Trial Day** | **Number of repetitions per each movement condition** |
| --- | --- | --- | --- | --- | --- |
| T5 | -4.5 | 11.15.2022 | 6, 7, 9, 10, 11, 12, 13, 14, 15, 16, 18 | 2281 | 22 |
| T11 | -4.5 | 09.27.2022 | 8, 9, 10, 11, 12, 13, 14 | 1099 | 14 |
| T12 | -4.5 | 04.19.2022 | 4, 5, 6, 7, 8, 9, 10, 11, 12, 13 | 21 | 20 |
| T15 | -4.5 | 08.27.2023 | 6, 7, 8, 9 | 41 | 8 |
| T16 | -3.5 | 01.10.2024 | 6, 8, 10, 11, 12, 13, 14 | 34 | 13 - 14 |
| T17 | -3.5 | 04.09.2024 | 1,2,3,4,5,6,7,9 | 40 | 16 |
| C1 | -4.5 | 06.23.2023 | 1, 2, 3, 4, 5, 6, 7, 8, 9, 10, 11, 12, 13, 14 | 969 | 14 |
| C2 | -4.5 | 06.19.2023 | 1, 2, 3, 4, 5, 6, 7, 8, 9, 10, 11, 12, 13 | 53 | 12 - 13 |

The cued movement task followed a simple instructed delay paradigm (Fig. 1b). During the instructed delay period a red square and text appeared in the center of the screen indicating to the participant that they should prepare to make the specified movement. The instructed delay period varied randomly (except for participants C1 and C2), with the range tailored to each participant’s preference for sufficient time to read the prompt and prepare the movement. After the delay, the square turned green and the text indicating the movement changed to “Go”, at which point the participant executed the movement immediately. The participant was instructed to make the movement if they were able to overtly move that body part, otherwise they were instructed to attempt the movement. They were then directed to continue attempting the movement, or holding the posture of the completed movement until the text changed to “Return”, at which point the participant relaxed and returned to a neutral posture. Typically, the movement period lasted 1.5 seconds and the return period 1 second, although these durations were adjusted if participants required more time to complete the task. See Table 2 for task timing parameters.

**Table 2.** Task timing parameters and analysis windows. The ‘Go’ period analysis window for each participant was defined as the entire ‘Go’ period while accounting for a 300 ms reaction time. The ‘Delay’ period window used for analysis was defined as a 1 s window prior to the ‘Go’ cue.

| Participant | Delay low | Delay mean | Delay high | Go period | Return period | Go period analysis window (relative to 'Go' cue) | Delay period analysis window (relative to 'Go' cue) |
| --- | --- | --- | --- | --- | --- | --- | --- |
| T5 | 1.5 s | 2 s | 2.5 s | 1.5 s | 1 s | 0.3 - 1.5 s | -1 - 0 s |
| T11 | 2 s | 2.5 s | 3 s | 1.5 s | 1 s | 0.3 - 1.5 s | -1 - 0 s |
| T12 | 2 s | 2.5 s | 3 s | 1 s | 1 s | 0.3 - 1 s | -1 - 0 s |
| T15 | 1.5 s | 2 s | 2.5 s | 2.5 s | 2.5 s | 0.3 - 2.5 s | -1 - 0 s |
| T16 | 2 s | 2.5 s | 3 s | 2.5 s | 1 s | 0.3 - 2.5 s | -1 - 0 s |
| T17 | 2.5 s | 3 s | 3.5 s | 2.5 s | 2 s | 0.3 - 2.5 s | -1 - 0 s |
| C1 | 3 s | 3 s | 3 s | 1.5 s | 2.5 s | 0.3 - 1.5 s | -1 - 0 s |
| C2 | 2 s | 2 s | 2 s | 1.5 s | 2.5 s | 0.3 - 1.5 s | -1 - 0 s |

## 2 Neural representation of whole-body movements in motor cortex

### 2.1 Peristimulus time histograms

To generate the peristimulus time histograms (PSTHs) shown in Fig. 1c, we first started with binned threshold crossing (TX) spike counts (20 ms bins). For visualization purposes, we denoised the data by convolving each electrode’s TX counts with a Gaussian smoothing kernel (120 ms s.d.). Next, for each electrode we extracted TX counts for each trial in a -2 s to 2.5 s time window relative to each trial’s ‘Go’ cue. For each of the movement conditions (see Fig. 1c legend), we computed the mean and 95% confidence intervals (CIs; estimated using Matlab’s *normfit* function) for the TX counts in each time bin across all trials. We then scaled the resulting means and CIs (multiplying by 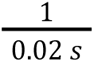) to convert them into units of Hz.

### 2.2 Neural tuning strength

Note that the neural population analyses discussed in this section rely heavily on the *cvVectorStats* library of functions for using cross-validation to estimate Euclidean distance between the means of multivariate distributions (and for estimating other statistics that require Euclidean distance, such as Pearson’s correlation). This library has been extensively used in our prior reports^11,18,40,64^ (see code repository https://github.com/fwillett/cvVectorStats).

#### 2.2.1 Tuning strength heat maps

Neural tuning strength heat maps shown in Fig. 2a were generated using cross-validated estimates of Euclidean distances between the distribution of population-level neural firing rates for a control ‘Do Nothing’ condition and the distribution of firing rates for a specific movement condition (see Fig. 2a column labels for each movement condition).

We first started with binned threshold-crossing (TX) counts (20 ms bins) for each array. To account for drifts in mean firing rates across the session, the TX rates were mean-subtracted within each block (i.e., for each electrode, its mean firing rate within each block was subtracted from each time step’s binned spike count). Next, for each trial and electrode, TX counts were averaged in the entire time window following the ‘Go’ cue, accounting for a 300 ms reaction time to cue onset (see Table 2 for analysis windows for each participant). This yields an Nx1 neural firing rate vector for each trial, where *N* is the number of electrodes. For each unique movement condition, including the ‘Do Nothing’ condition, we stacked each trial’s firing vectors to create a TxN matrix, where *T* is the number of trials for that specific movement condition. Next, we estimated the Euclidean distance between the ‘Do Nothing’ firing vectors and each unique movement condition’s firing vectors using the *cvDistance* function in the cvVectorStats library*. cvDistance* also returns 95% confidence intervals via jack-knife resampling (see Table 3 for statistical details). If the confidence interval contained 0, the tuning strength was considered insignificant relative to the ‘Do Nothing’ condition (denoted by a white ‘X’). To compare tuning strengths across arrays and participants whose neural data may differ in signal quality, the Euclidean distances calculated for each array (i.e., each row of the heat map) were normalized by the maximum Euclidean distance in that row.

**Table 3.** Statistical details.

|  |  |
| --- | --- |
| Figure 1 | Mean firing rates for each movement condition in Figure 1c were taken over 22 trials for T5, 8 trials for T15, 20 trials for T12, and 13 or 14 trials for T16. 95% confidence intervals were estimated using Matlab's *normfit* function which fits a normal distribution to the sample data. |
| Figure 2 | Statistical significance of tuning strength in panel A (each cell of the heatmap prior to row-wise normalization) was assessed by whether the 95% confidence interval intersected 0. See Table 1 for the number of trials for each movement condition that was used to compute the euclidean distance and confidence intervals. 95% confidence intervals were computed via jack-knife resampling (10,000 resamplings). See *cvVectorStats* package *cvDistance*.<br>There was no test for significance in panel B. Set-wise average tuning strengths were computed by averaging over the tuning strengths for the movements contained within each set: 4 movements within the Speech set, 5 movements within the Face set, 4 movements within the Head set (2 for C1), 8 movements within the Right Arm set, 8 movements within the Right Leg set, 8 movements within the Left Arm set, and 8 movements within the Left Leg set. |
| Figure 3 | 5-fold cross-validation was used for the discrete classification results shown in Fig. 3a. Significant decoding accuracy was assessed by computing the 95% confidence interval via the Clopper-Pearson method (Matlab's *binofit*). Chance level was computed for each movement set as 1/(number of conditions). If the confidence interval enveloped the chance level, it was deemed non-significant decoding performance. See Table 1 for the number of samples (trials) per condition. |
| Figure 5, Extended Data Figure 5 | Each correlation matrix in Figure 5a is the average across the correlation matrices computed for each array in the set (10 correlation matrices for 'Dorsal PCG', and 2 matrices for superior-ventral PCG')<br>Statistical significance in Fig. 5b was assessed via a shuffle-control to demonstrate that the movement correlations are significantly above chance. The correlation matrix for a given movement pair (column) for an array (row) was reshuffled (10,000 reshuffles of the rows and columns) and the mean of the diagonal entries were computed for each reshuffle. The 2.5th and 97.5th percentiles were computed on the sample means and served as the lower and upper limits of the 95% bootstrap interval for the shuffle control. If the mean diagonal entries of the original correlation matrix were outside of the 95% bootstrap interval, it was deemed significant.<br>The number of trials in each 2D projection for Figure 5c and Extended Data Figure 5 is:<br>T5 : 176 Right arm trials, 176 Left arm trials<br>T11 : 112 Right arm trials, 112 Left arm trials<br>T12 : 160 Right arm trials, 160 Left arm trials<br>C1 : 112 Right arm trials, 112 Left arm trials<br>C2 : 104 Right arm trials, 104 Left arm trials<br>T15 : 64 Right arm trials, 64 Left arm trials<br>T16 : 112 Right arm trials, 112 Left arm trials<br>T17 : 128 Right arm trials, 128 Left arm trials |
| Extended Data Figure 1 | Significant tuning to a movement condition for each electrode in Panel A was assessed via a 2-sample t-test (Matlab *ttest2* function). The number of samples (trials) per group is shown in Table 1. If the p-value was less than 1e-5, it was deemed significant. |
| Extended Data Figure 7 | 10-fold cross-validation was used for the discrete classification sweeps. Statistical significance in decoding performance was assessed via computing the 95% confidence interval via the Clopper-Pearson method (Matlab's *binofit*). Chance level was computed as 1/(total movement conditions). If the confidence interval intersected chance level, decoding performance was deemed non-significant. See table 1 for the number of samples (trials) per condition for each participant. |

#### 2.2.2 Movement set-wise tuning strength across spatially organized arrays

For Fig. 2b, we plotted the microelectrode arrays as they appear on PCG to better interpret how they are tuned as a function of their spatial location. We used anatomical landmarks to position the arrays relative to each other across participants in a generic space.

The brain surface depicted in the left-most panel of Fig. 2b is the Human Connectome Project’s group-averaged pial surface (HCP S1200^16,65^). The superior frontal sulcus (SFS) and inferior frontal sulcus (IFS) are illustrated by hand-drawn black lines and are easily identifiable landmarks despite variability in brain folding across individuals. We then defined the extremes of the PCG surface as ‘Superior’ and ‘Inferior’. Using the MRI-derived brain anatomies for each participant, we approximated each array’s location relative to these four landmarks for plotting purposes (right panels of Fig. 2b).

For each microelectrode array, tuning strength to each movement set (Speech, Face, Head, Right Arm, Right Leg, Left Arm, Left Leg) was computed by averaging the tuning strengths across the sub-movements within each set (i.e., averaging the relevant entries in Fig. 2a before row-wise normalization). Sub-movements with no statistically significant tuning (i.e., cells with white ‘X’s) were still included in the average. Computing set-wise averages yields a tuning strength matrix of size AxS, where *A* is the number of arrays and *S* is the number of movement sets. The set-wise tuning strength matrix was then row-wise normalized by the maximum value. Next, each microelectrode array (square) in Fig. 2b was colored according to the corresponding value in this normalized matrix.

### 2.3 Recurrent neural network classifier

For Fig. 3, we used the recurrent neural network decoding (RNN) architecture described in our prior work^40^ to classify movements using single trial neural recordings. For each trial, binned TX spike count time series from a four second window after the go period were used as input to the RNN. Binned TX spike counts were block-wise mean subtracted to remove firing rate drifts across the session. The RNN was trained on all movement classes and arrays simultaneously, with a separate input layer for each array (using the same methods as in^40^, where separate input layers were used for each day of data). Training separate input layers helps to place the data into a common space before processing by the RNN. To account for different array sizes (64 vs 96 electrodes), data from 64-electrode arrays was zero-padded to increase the dimensionality to 96. All hyperparameters were the same as that used in our prior work^40^, except instead of using the CTC loss, we used a cross-entropy loss applied to a linear readout from the last RNN layer (using the PyTorch function *torch.nn.CrossEntropyLoss*).

We used 5-fold cross-validation to estimate classification accuracy (5 RNNs were trained on 80% of the trials each and tested on the remaining 20%). Only a single RNN was trained for each fold to classify *all* movements across *all* arrays (as opposed to training RNNs separately for each movement category and array). When computing classification accuracy for a particular movement category, the decoder output was constrained to be one of the relevant movements from that category only (the maximum probability output within the allowable set was chosen as the final output). Classification accuracies were considered statistically significant if their 95% confidence intervals did not intersect chance level. 95% confidence intervals for the classification accuracies were calculated using Matlab’s *binofit.* The chance level was calculated as 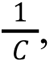 where *C* is the number of conditions within each set.

### 2.4 Principal component analysis of array tuning properties

To make Fig. 4a, normalized modulation magnitudes for each category of movement (as depicted in Fig. 2a), and classification accuracies (as depicted in Fig. 3a) were concatenated together for each array to produce a 20 x 14 matrix (20 arrays, 7 + 7 accuracies and normalized modulation magnitudes). Classification accuracies were expressed as success rates varying from 0 to 1, and normalized modulation magnitudes also varied from 0 to 1. Principal components analysis was then applied to the rows of this matrix to find the dimensions that explained the most variance in array tuning properties (these two principal components are depicted in Fig. 4b). Array tuning properties were then visualized in this two-dimensional space (Fig. 4a), and each array was colored using an HSV color scheme. The hue of each point (*x*, *y*) in this space was determined by the angle *atan*2(*y*, *x*), the saturation was determined by the magnitude 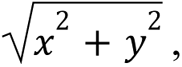, and the value was set to 0.85. Saturation values were clipped at 1.

### 2.5 Correlation between homologous limb movement representations

Fig. 5a and Extended Data Figure 4 show average pairwise correlations (Pearson’s r) between the neural representations of homologous limb movements.

First, following the same pre-processing steps as section 2.2.1, binned TX spike counts for each microelectrode array were block-wise mean removed, and neural firing rate vectors were computed by averaging the TX counts within the ‘Go’ period window (see Table 2) for each trial. Firing rate vectors were then concatenated into TxN matrices for each movement condition *m*, where *T* is the number of trials for movement *m* and *N* is the number of electrodes within an array. Before computing the correlations, we subtracted the average firing rate within each movement set to remove any effector-dependent neural dimensions such as the potentially large laterality and arm versus leg dimensions reported in prior work^11,18^. For example, for arm movements, we averaged across all neural vectors for movements within the arm set, creating a 1xN average vector which was then subtracted from the neural vectors within this arm set. Next, for each pair of movements (*m*_i_, *m*_j_), the corresponding concatenated firing rate matrices were used to compute the cross-validated correlation between the mean firing rate vectors of each movement (*cvCorr* function from the cvVectorStats library). The correlation matrices were then averaged across arrays within the following sets: dorsal PCG, middle PCG, superior-ventral PCG, and inferior-ventral PCG. See the row labels of Fig. 2a for how the arrays were grouped into these sets.

For Extended Data Figure 4, note that the row-column entries of the correlation matrices for ‘Right arm -Left arm’ and ‘Right leg - Left leg’ were re-ordered to align movements that are the same in joint-angle space as opposed to extrinsic, Cartesian space. For example, ‘Right arm - Raise right is homologous to ‘Left arm - Raise left’. This aligns with our previous reports which showed that directional movements were correlated across effectors in intrinsic space^11,18^.

The heat map in Fig. 5b indicates the average correlation between homologous movements for limb pairs (column entries) for each microelectrode array (row entries). Each cell represents the mean of the diagonal values of the corresponding correlation matrix for each limb pair. A statistical test for significance was performed by a shuffle-control test (black ‘X’ indicates non-significant correlation between homologous movements). See Table 3 for statistical details.

### 2.6 Movement-independent neural coding of laterality

We used PCA to visualize the neural activity, in select microelectrode arrays, in a lower-dimensional space as illustrated in Fig. 5c. First, binned TX spike counts were block-wise mean removed and z-scored (mean-subtracted and divided by the standard deviation) for visualization purposes. Next, firing rate vectors were computed for each trial by averaging the counts within the ‘Go’ period windows (see Table 2). We then concatenated all of the firing rate vectors for right and left arm movements into a matrix of size TxN, where *T* is the total number of trials and *N* is the number of electrodes. PCA was performed on this monolithic matrix and each z-scored firing rate vector was subsequently projected onto the top two principal components (PCs). The single-trial projections were colored by the corresponding movement set (right arm trials in red, left arm trials in blue). See Table 3 for details on the number of trials.

The heat map in Fig. 5d summarizes the size of laterality-related tuning using a variation of demixed principal component analysis^66^ (dPCA; Kobak et al., 2016; https://github.com/machenslab/dPCA). A core concept of dPCA involves marginalizing neural data across different experimentally manipulated factors, or variables. Each marginalization averages across variables not in the set, creating a data tensor which captures the effect of those factors on the neural activity. Using the dPCA library, we applied a cross-validated variance computation to estimate the amount of variance in the neural activity due to each factor while reducing bias^18^. To reduce bias when estimating variance, we split the trials into two folds and computed the marginalizations separately for each fold. We then estimated each marginalization’s covariance matrix as *X*_1_ *X*_2_^*T*^, where *X* is the marginalization computed on fold 1 and *X* is the marginalization computed on fold 2. We then took the real part of the eigenvalues of *X*_1_ *X*_2_^*T*^ to estimate each component’s variance. The data was marginalized into the following four factors: laterality (left or right arm), movement type (Arm - Raise left, Arm - Raise right, Hand - close, Hand - open, Wrist - Up, Wrist - Down, Wrist - Left, Wrist - Right), laterality and movement type interaction, and time. We computed the cross-validated variance in the aforementioned factors for each microelectrode array. Each cell of the heat map in Fig. 5d represents the cross-validated marginalized variance for each factor (labeled along the x-axis) across each microelectrode array.

### 2.7 Intermixed tuning within electrodes

Extended Data Figure 1a shows matrices which indicate whether an electrode was significantly tuned to an individual movement. To assess significant tuning, we first started with binned TX spike counts which were block-wise mean removed. Next, for each trial and electrode, we averaged the TX counts within the ‘Go’ period window. Significance of tuning was then assessed via a two-sample t-test (using the *ttest2* function in Matlab) applied per electrode, where group 1 was comprised of the single-trial average firing rates for movement *m*, and group 2 was comprised of the single-trial average firing rates for the ‘Do Nothing’ condition (see Table 1 for number of trials per condition). This two-sample t-test was performed for each movement (p < 0.00001 defined significance). Electrodes which were significantly tuned to a movement appear as a filled in circle in Extended Data Figure 1a.

The heat map in Extended Data Figure 1b summarizes the fraction of electrodes that exhibited significant tuning to at least one movement within each movement set as computed in Extended Data Figure 1a. The heat map in Extended Data Figure 1c summarizes the fraction of electrodes that had statistically significant tuning to each possible number of movement sets (from 0 to 7). The color bar is clipped at 0.5 to better visualize the range of values.

### 2.8 Data exclusion

Microelectrode array recordings may not be representative of the neural population tuning in a cortical area if they fail to record a sufficient amount of tuned spiking activity. To test whether an array contained any movement-related information, we used a cross-validated (10-fold) Gaussian naive Bayes classifier (following the methods described in^11^) applied to a 100 ms sliding window of neural activity. We found that 4 microelectrode arrays (C2-d_1_, T11-d_2_, T17-m_1_ and T17-m) failed to demonstrate consistently above-chance classification performance for any time epoch, when classifying from amongst all 46 movements (Extended Data Figure 7), and were thus excluded from this study and all other main results.

Also of note, is that participant C1’s neural recordings exhibited large noise when they attempted movements of the head moving up and down, which may have mechanically disturbed the pedestals fixed to the head or the cables. These two head conditions were removed for participant C1 for all population-level tuning analyses.

## 3 Human Connectome Project parcellation, resting state networks, and motor task activations

For all Human Connectome Project (HCP) results presented in this study, we analyzed the WU-Minn HCP 1200 Subjects Group Average Data Release^16,65^ and the original 210 subject release^16^. This data includes group average structural data, functional connectivity data, and task fMRI data. The Connectome Workbench (v1.5.0; www.humanconnectome.org), an open-source visualization and discovery tool, was used to explore and analyze the data generated by the HCP.

### 3.1 Group cortical parcellation

Extended Data Figure 3 (first panel) displays the group-averaged pial surface of the left hemisphere with 180 areas delineated and identified by the HCP’s multi-modal parcellation^16^ (HCP_MMP1.0). To generate this panel, we loaded the HCP_S1200_GroupAvg_v1.scene file into the Connectome Workbench and selected the ‘Cortical Parcellations’ scene (scene 5). Next, we viewed the left hemisphere’s pial surface (S1200.L.pial_MSMAll.32k_fs_LR.surf.gii) and overlaid the cortical parcellations (Q1-Q6_RelatedValidation210.CorticalAreas_dil_Final_Final_Areas_Group_Colors.32k_fs_LR. dlabel.nii) with modifications to the color scheme. We enabled the borders and selected only the following regions to display: L_4_ROI, L_6mp_ROI, L_6d_ROI, L_6v_ROI, L_PEF_ROI, L_FEF_ROI, L_55b_ROI.

### 3.2 Resting state fMRI networks for language and arm movement

The resting state networks shown in Extended Data Figure 2 were generated using the same scene file in 3.1 and overlaying resting state networks 25 and 28 (Q1-Q6_RelatedParcellation210.individual_RSNs_d40_WR_norm_MSMAll_2_d41_WRN_De Drift.32k_fs_LR.dscalar.nii; file can be found at https://balsa.wustl.edu/study/RVVG). Networks 25 and 28 were highlighted as a language network and upper limb network, respectively, by Glasser et al. 2016^16^.

### 3.3 Task fMRI activations

Extended Data Figure 6 shows task fMRI activations for the group averaged HCP S1200 dataset. The same main scene file from section 3.1 was used with the ‘S1200 task fMRI Cohen’s D effect-size maps’ scene loaded (scene 3). Task activation file HCP_S1200_997_tfMRI_ALLTASKS_level2_cohensd_hp200_s2_MSMAll.dscalar.nii was used for overlay visualization and analysis.

Extended Data Figure 6a shows sulcal depth maps (S1200.sulc_MSMAll.32k_fs_LR.dscalar.nii) on the inflated group-averaged brain surface (S1200.L.inflated_MSMAll.32k_fs_LR.surf.gii). The inflated brain surface was used to more easily select probing points. Black lines indicate areal borders for the parcellated areas. Within the Connectome Workbench, we hand selected points (white dots) on the brain surface along the fundus of the central sulcus distinguished by deeper sulcal depth (following the dark blue regions), and along the crest of PCG (following the dark red regions). For each point, we extracted the vertex value to compute the distance between the probed points, and the associated row-index into the task activation niftii file to extract the activation value for each tested movement. The task fMRI movements are left foot, left hand, right foot, right hand, and tongue (column-indices 38, 39, 40, 41, 42 of the niftii file, respectively). The cumulative distance between probed points is plotted against the task activations extracted from the niftii file in Extended Data Figure 6b.

## 4 Statistics

Table 3 lists statistical details for each confidence interval or hypothesis test reported in this work. In this study, uncertainty was quantified mainly with 95% confidence intervals.

